# The dynamic genomes of Hydra and the anciently active repeat complement of animal chromosomes

**DOI:** 10.1101/2024.03.13.584568

**Authors:** Koto Kon-Nanjo, Tetsuo Kon, Tracy Chih-Ting Koubkova Yu, Diego Rodriguez-Terrones, Francisco Falcon, Daniel E. Martínez, Robert E. Steele, Elly Margaret Tanaka, Thomas W. Holstein, Oleg Simakov

## Abstract

Many animal genomes are characterized by highly conserved chromosomal homologies that pre-date the ancient origin of this clade. Despite such conservation, the evolutionary forces behind the retention, expansion, and contraction of chromosomal elements, and the resulting macro-evolutionary implications, are unknown. Here we present a comprehensive stem-cell resolved genomic and transcriptomic study of the fresh-water cnidarian *Hydra*, an animal characterized by its high regenerative capacity, the ability to propagate clonally, and an apparent lack of aging. Using single-haplotype telomere-to-telomere genome assemblies of two recently diverged hydra strains, we show how the macro-evolutionary history of chromosomal elements is shaped by both old and recent transposable element (TE) expansions. Unique features of hydra biology allowed us to compare the individual genomes of hydra’s three stem cell lineages. We show that distinct TE families are active at both transcriptional and genomic levels via non-random insertions in the genomes of each of these lineages. In transcriptomes, over 14,000 transcripts were composed of nearly complete TE sequences, and further classification into families, subfamilies, and individual loci reveals cell type-specific TE expression. The active TEs include elements that differentially contribute to changes in the genome size as well as persistent structural variants around loci associated with cell proliferation. Our study reveals 14 active TE families that primarily act in this role and are predominantly composed of DNA elements. Evolutionary analysis revealed that these families constitute a highly conserved TE core in eukaryotic and metazoan genomes. Our results suggest an ancient role for these core TEs as self-renewing genomic components that persist beyond ancient chromosomal homologies.

## Introduction

Animal genomes have been shown to be remarkably conserved at both chromosomal^1,2^ and sub-chromosomal levels^3–5^. While this remarkable conservation can be explained by gamete balancing issues that can arise in the case of inter-chromosomal translocations in the germline^6^, the evolutionary forces that keep regulatory landscapes intact within chromosomes are less understood.

One such force is the continuous expansion and contraction of chromosomes through the action of transposable elements (TEs). How this dynamic chromosomal landscape may contribute to macroevolution, and the resulting evolutionary constraints on chromosomal elements themselves is unclear. Despite the recent increase in the availability of chromosomal-scale genome assemblies for many animal species, high-resolution comparative genomics of TEs requires telomere-to-telomere haplotype-resolved genome assemblies and a detailed understanding of TE activity and insertion dynamics for closely related species. Very few such resources are available. The freshwater cnidarian *Hydra* is a powerful system for addressing these questions. As a member of the sister phylum to bilaterians, hydra occupies an important evolutionary position. Hydra was the first animal in which whole body regeneration was demonstrated^7^, and the first animal in which a developmental organizer was identified^8,9^. Hydra serves as a simple yet powerful model system, providing profound insights into stem cell dynamics. The adult hydra polyp does not age^10^. Homeostasis and the extensive regeneration capacity in hydra are maintained by three stem cell lineages, endodermal epithelial, ectodermal epithelial, and interstitial/germline stem cells (i-cells). Endodermal and ectodermal epithelial stem cells are unipotent, only producing epithelial cells. I-cell stem cells are multipotent, and produce nematocytes, nerve cells, gland cells, and germ line cells^11^, though the rate may depend on the developmental stage^12^. The three stem cell lineages maintain their distinct identities and do not interconvert. Hydra regeneration from bisected animals, small tissue fragments, and even aggregates of cells has been extensively studied since the first experiments by Abraham Trembley in the 1700s^7^. More recently, transcriptomic studies have revealed the diversity of the molecular pathways that are active during hydra regeneration^13^. Comprehensive profiling of gene expression in each cell type and stem cell differentiation trajectories have been reported for the whole polyp of hydra through single cell RNA-seq (scRNA-seq)^14^. While components such as the conserved Wnt signaling pathway have been investigated in developmental and regeneration process in hydra, many species-specific genes and, particularly, TEs are expressed, with unclear functional or evolutionary impact^13^.

The ability of hydra to propagate asexually has striking implications for its biology. If hydras are cultured by asexual budding for multiple generations, mutations accumulate in each stem cell lineage independently. As a consequence, clonally propagated hydras are expected to have three distinct genomes (ectodermal, endodermal and i-cell stem cell lineages). If TEs have diverging functions in each of the three genomes, we expect to find differences in their cell-type specific transcriptional activity and genomic insertion profiles. These profiles may suggest potential impacts on the cellular function and adaptation.

The genus *Hydra* is composed of two major evolutionary lineages, the symbiotic green hydra (*H. viridissima*) and the species-rich brown hydras (e.g. *H. vulgaris* and *H. oligactis*)^15^. These two lineages diverged 61–46 million years ago^15^, yet in this relatively short evolutionary time-span, brown hydra genomes have experienced a tripling in genome size^16,17^ while still preserving the ancient animal chromosomal homologies^1^. The green hydra genome is similar in size to genomes of most other cnidarians (less than 500 Mbp)^18^, the brown hydras have some of the largest cnidarian genomes (around 1 Gbp or more), with over 60% of the total sequence comprised of repetitive sequences, including TEs^16,19^.

Within the brown hydras, two strains, AEP and 105, have been extensively utilized in the study of hydra biology. The AEP strain of *H. vulgaris*, from North America^20,21^, routinely undergoes sexual reproduction in laboratory conditions and is widely used to create transgenic lines^22^. Conversely, the 105 strain of *H. vulgaris*, which was collected in Japan in 1973, has been maintained through asexual reproduction in the laboratory^23^. Although it almost certainly engaged in sexual reproduction in its natural environment, it has ceased undergoing sexual reproduction since its establishment as a laboratory strain.

To uncover the role of the evolving TE landscape in hydra stem cell genome(s) and its impact on shaping the otherwise preserved, yet expanded, ancient animal chromosomal elements, we present the haplotype-resolved telomere-to-telomere assemblies of *Hydra vulgaris* 105 and AEP strains and reveal different TE evolutionary expansions among those lineages. Using fluorescence-activated cell sorting (FACS) followed by genome sequencing we reveal distinct TE insertion profiles in the three stem cell lineages in the AEP strain. We show evidence indicating how these TE families impact the genome and target genes. We provide the first direct evidence that the three stem cell genomes of hydra undergo distinct and non-random changes and suggest this as a key structural role of TEs in the evolutionary adaptations of these stem cells. Finally, through multi-species comparisons, we find remarkably similar TE signatures across eukaryotes and metazoans, identifying core sets of TE elements that have shaped chromosomal evolution for billions of years.

## Results

### Haplotype-resolved telomere-to-telomere reference assemblies for 105 and AEP strains of *H. vulgaris*

While chromosomal-scale assemblies of *H. vulgaris* strain 105 and strain AEP have been published^1,19^, they lacked telomere-to-telomere completeness and haplotype resolution needed to study the action of TEs in a stem cell lineage resolved manner. We utilized newly generated chromosomal conformational capture data from a AEPx105 F1 hybrid (Methods) and new long-read sequencing to generate telomere-to-telomere assemblies (Fig. 1a,b, Supplementary Fig. 1a–f, Supplementary Data 1–3). As the sequencing data came from a hybrid of two recently diverged hydra strains (Fig. 1a, Supplementary Fig. 1a), our data produced a single haplotype for each of the strains with telomere-to-telomere quality (Fig. 1a,b, Supplementary Fig. 1b–f, 2a,b). Both strains showed a high degree of local (micro-) synteny conservation and the previously reported conservation of ancient animal chromosomal elements (Fig. 1c)^1^. However, the 105 strain genome showed a single apomorphic translocation between chromosome 5 and chromosome 15, which was previously suggested^1^ and can now be confirmed with our data (Fig. 1c). Our analysis further identifies this event as a Robertsonian translocation where chromosome arms are exchanged at the centromeric regions (Fig. 1c, Supplementary Fig. 2c–e).

**Fig. 1.**
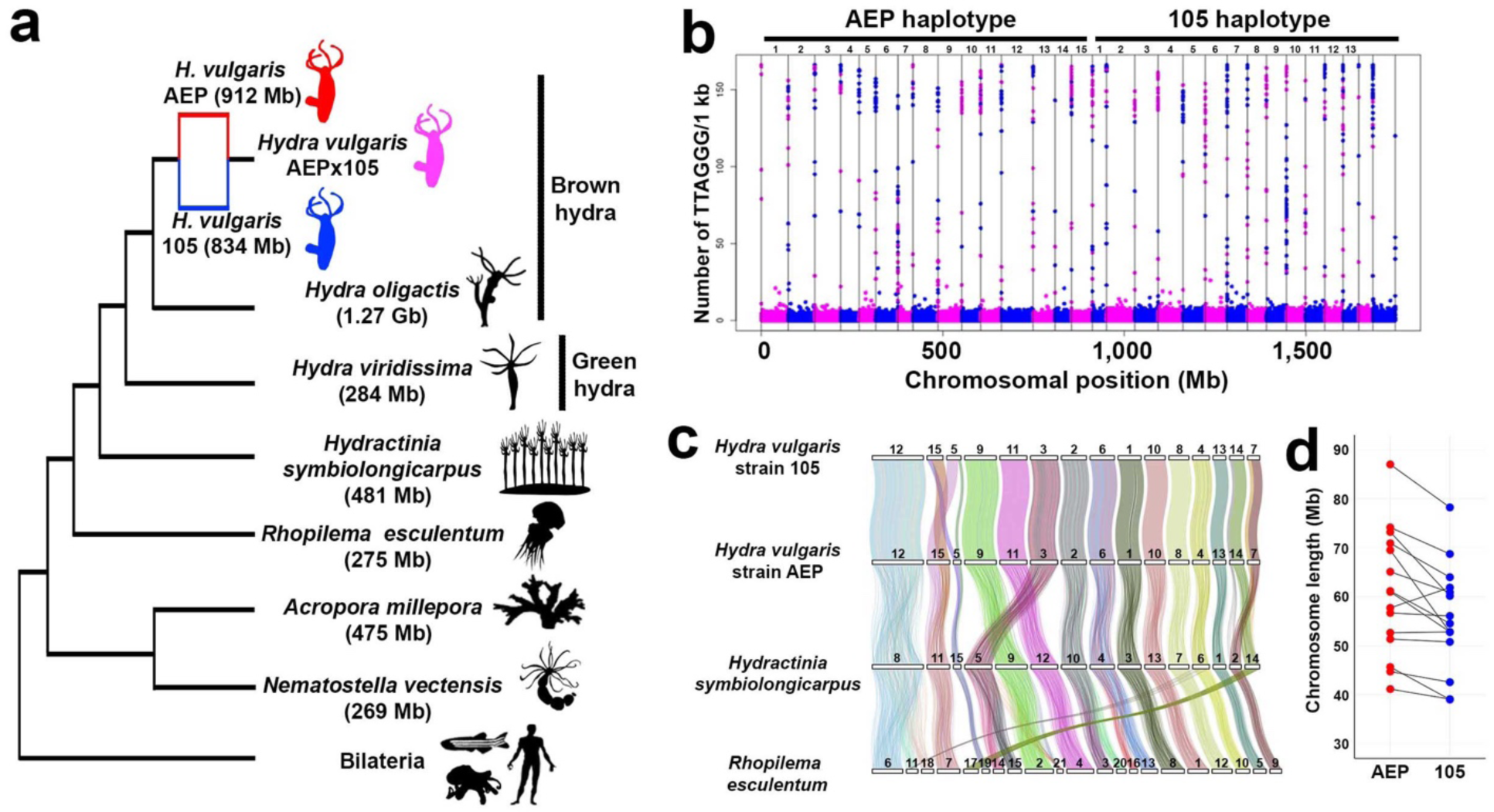
Haplotype-resolved telomere-to-telomere genome assemblies for 105 and AEP strains of *H. vulgaris*. **(a)** Cladogram representing major cnidarian species. Numbers in parentheses indicate the sizes of publicly-available genome assemblies. Animal silhouettes are from PhyloPic (www.phylopic.org). **(b)** Distribution of telomeric repeats. The x-axis represents genomic positions and the y-axis shows the number of occurrences of telomeric repeats (TTAGGG) per kilobase pair. The plots of each chromosome are colored alternately to distinguish between the plots of the terminal regions of two different chromosomes. **(c)** Deeply conserved synteny of chromosomes of *Hydra vulgaris*, *Hydractinia symbiolongicarpus*, and *Rhopilema esculenta*. The coordinates of 3,685 orthologs that form 27 ancestral linkage groups are connected by lines. Each line color corresponds to one of the 27 ancestral lineage groups. **(d)** Chromosome lengths of each haplotype genome. Plots between orthologous chromosomes are linked with black lines.

When comparing the 13 chromosomes, excluding the chromosomes that underwent the Robertsonian translocation, between each haplotype genome, 12 chromosomes of the AEP haplotype were longer than those of the 105 haplotype suggesting that the AEP genome has experienced a genome-wide size increase (Fig. 1d). It was found that the major groups of TEs (including SINE, LINE, LTR, DNA, RC; Fig. 2a) accounted for 65.3% (596 Mb) of the AEP genome and 63.3% (528 Mb) of the 105 genome (Fig. 2b,c). Among these groups, DNA transposons were the most common, comprising 34.2% and 30.1%, followed by LINEs at 16.7% and 16.3% in the AEP and 105 genomes, respectively (Fig. 2b). When comparing the age distribution (Kimura substitution levels) among each major TE group between the AEP and the 105 haplotypes, we found that relatively young DNA elements and LINEs have increased in AEP, suggesting their contribution to the genome size increase after the divergence of the last common ancestors of AEP and 105 (Fig. 2d).

**Fig. 2.**
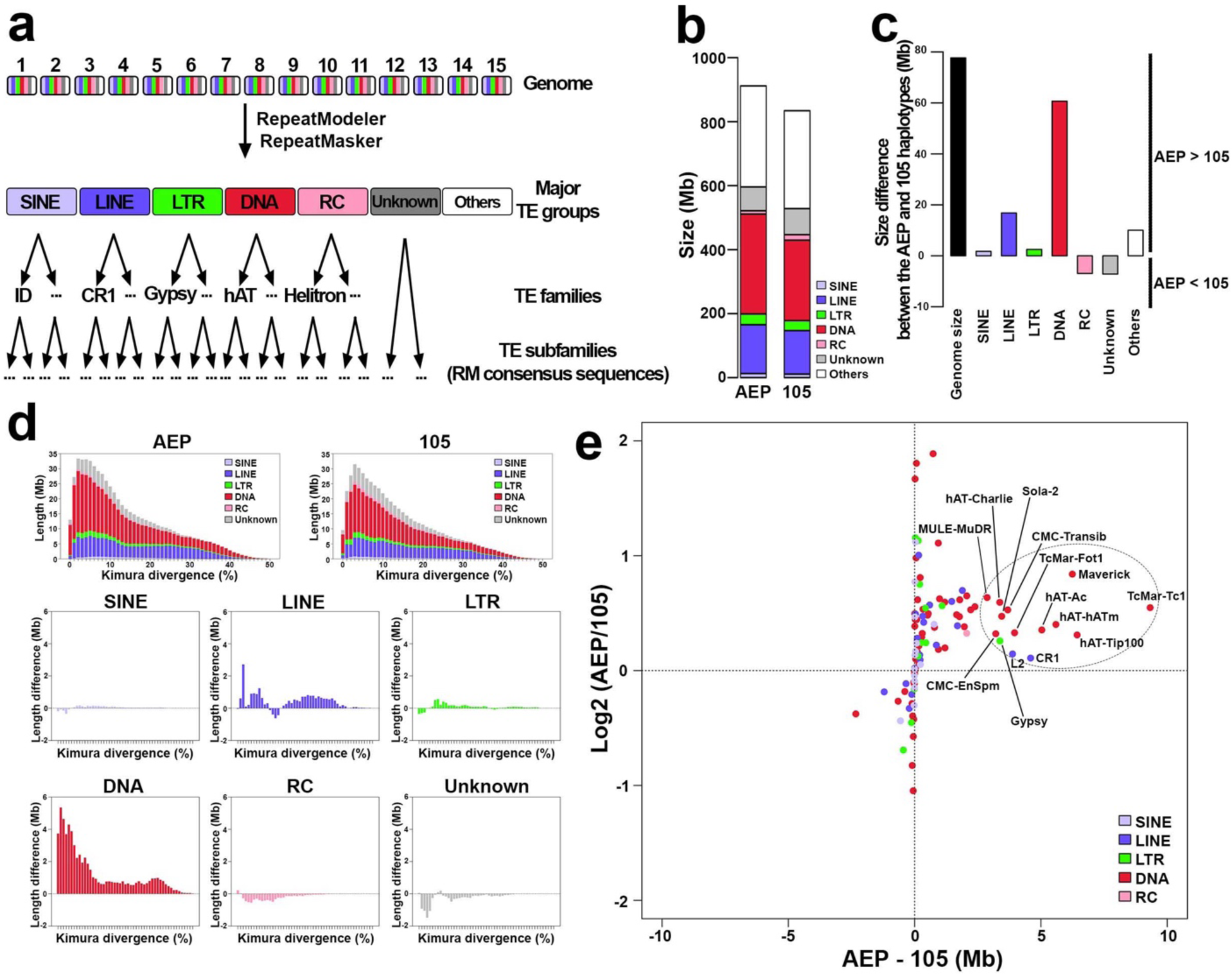
DNA transposons as the major contributor behind the recent AEP genome expansion. **(a)** TE classification pipeline and levels of TE annotation, RM: RepeatMasker. **(b-c)** Repetitive element content and genome size differences, and individual TE class contribution to 105 and AEP *Hydra vulgaris* strains. **(d)** Difference of Kimura divergence between the two strains showing enrichment and recent insertions of mostly DNA, but also LINE and LTR elements in the AEP strain. **(e)** Size differences and fold-changes of TE families between the AEP and 105 genomes highlighting specific TE family enrichment. The dotted eclipse highlights a cluster composed of 14 TE families with high contribution to the AEP genome size increase (A-TEs).

We used these genome assemblies to compare the two strains in detail and to identify changes along their chromosomes. The total size difference of TEs between the AEP and 105 haplotype genome is 67.6 Mb, which accounts for 87.1% of the difference in the genome size (Fig. 2c). In particular, we identified the putatively active TEs (“A-TEs”) composed of 14 TE families with the enrichment in AEP including 11 DNA elements (hAT-Ac, hAT-Charlie, hAT-hATm, hAT-Tip100, TcMar-Tc1, TcMar-Fot1, Maverick, Sola-2, MULE-MuDR, CMC-EnSpm, and CMC-Transib), two LINE elements (CR1 and L2), and one LTR element (Gypsy). These elements were found to contribute the most to the AEP genome size increase (Fig. 2e, Supplementary Data 4). Subfamilies of each family from these A-TEs show an even more diverse size distribution between the AEP and 105 genomes. For example, only few TcMar-Tc1 and CR1 subfamilies were enriched in the AEP genome, but the majority of hAT-Ac and Gypsy subfamilies had a contribution to the AEP genome size increase (Supplementary Fig. 3). While the 14 TE families of the A-TEs collectively affect whole chromosomes, some subfamilies of A-TEs showed local propagation in specific chromosomal regions in AEP characterized by the lack of local synteny, suggesting expansion through tandem duplication (Supplementary Fig. 4a,b).

This analysis, based on complete and haplotype-resolved assemblies, highlights a diverse set of putatively active TE (A-TE) families in the hydra genome, which suggests a more complex picture of the brown hydra genome expansion than the previous finding of a single retroelement contribution to the genome size increase in this clade^16,17^. We next sought to investigate these A-TEs transcriptionally.

### Expression activity of transposable elements in hydra

While expression of TEs has been proposed in hydra in previous studies^13,24^, validation using different sequencing technologies has been lacking. To identify the set of TEs that are substantially expressed, we analyzed an Iso-Seq transcriptome generated from whole polyps of the 105 x AEP hybrid line. Out of the 9,434,059 Iso-Seq reads, we searched for those containing more than 1 kb of TE-annotated sequences and found that 29,833 transcripts (0.32%) - 14,302 transcripts (0.15%) from the AEP haplotype genome and 15,531 transcripts (0.16%) from the 105 haplotype genome - were derived from TEs. These transcripts include the major groups of TEs: SINE, LINE, LTR, DNA, and RC (Fig. 3a,b), and encompass 86 different TE families (75 families and 73 families for the AEP and 105 haplotype genomes, respectively) (Fig. 3c). The average transcript length suggests that near full-length transposon sequences are being captured by Iso-Seq, with the shortest being 1.8 kb representing a SINE from the AEP haplotype genome and the longest being 3.5 kb containing an LTR from the 105 haplotype genome (Fig. 3a). We also find that 30.9% of TE Iso-Seq reads in AEP and 29.7% of TE Iso-seq reads in the 105 strain have trans-spliced leader (SL) sequences^16,25^ at the 5’ end of the reads, while for Iso-Seq reads of non-TEs, 48.1% were found to have SL sequences, suggesting that transcripts of TEs are utilizing hydra’s endogenous trans-splicing mechanism (Fig. 3b, Supplementary Fig. 5a), similar to the protein-coding genes^16^. All previously identified putatively active TEs (A-TEs) showed expression (Fig. 3c). Of all A-TEs, 11 A-TEs, excluding CMC-Transib, Maverick, and MULE-MuDR, were among the top 20% of the 86 TE types according to the expression level (Fig. 3c). All A-TEs showed high expression support (Fig. 3c). The TE-derived sequences were mapped to 4,672 loci in the AEP haplotype genome and to 4,087 loci in the 105 haplotype genome (Fig. 3d). At the subfamily level, expression of 868 subfamilies and 762 subfamilies were observed from the AEP and the 105 haplotype, respectively.

**Fig. 3.**
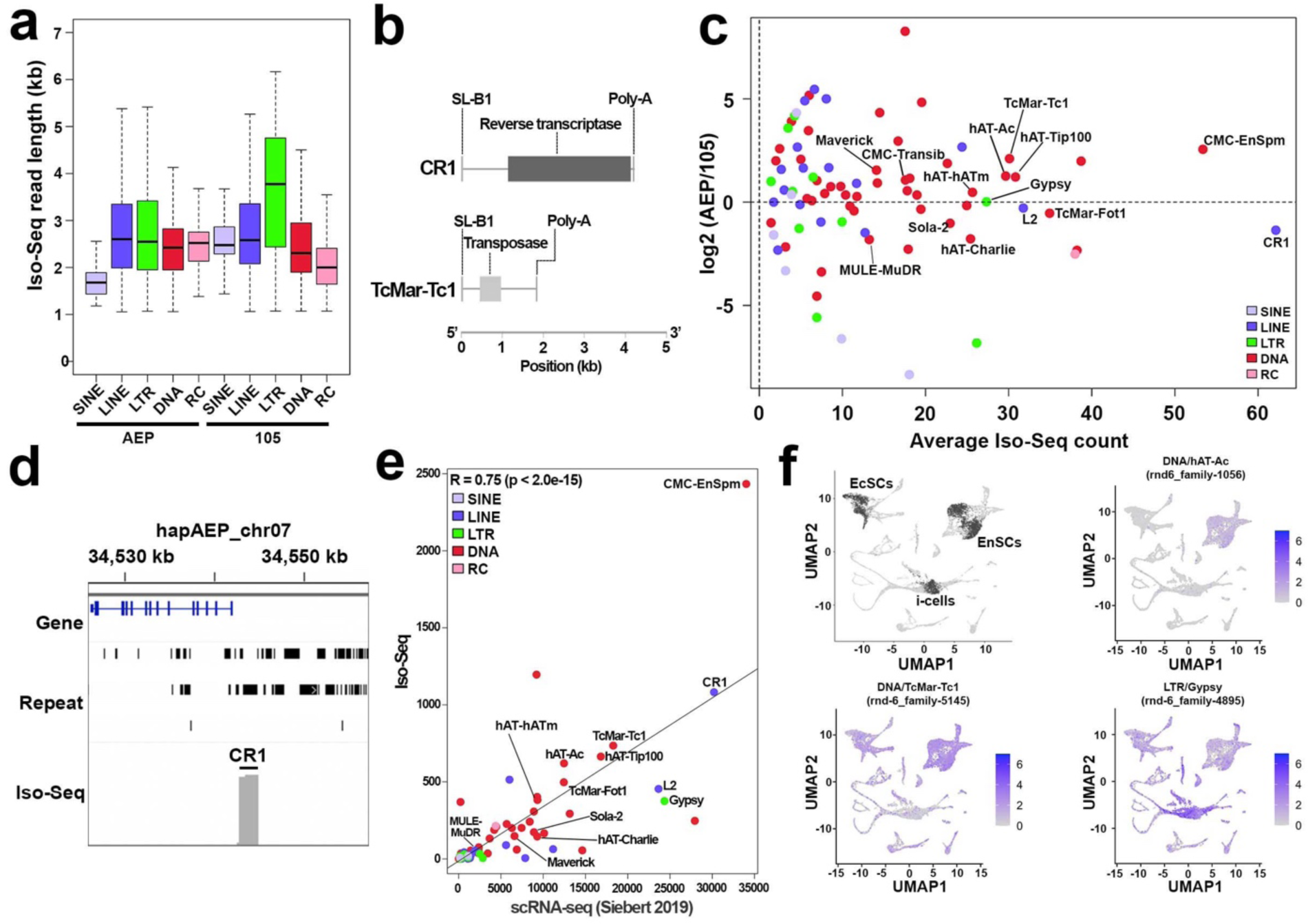
Strain and cell-type specific expression dynamics of Hydra TE families. **(a)** The lengths of Iso-Seq reads capturing TE sequences. The boundaries of the box depict the first (25%) and third (75%) quartiles, while the median is denoted by the central horizontal line within the box. The whiskers extending from the box indicate the lowest and highest observed values. **(b)** Full length transcripts of the CR1 element and TcMar-Tc1 element with trans-spliced leader sequences (SL-B1^16^) attached at the 5’ end**. (c)** Total expression difference (log2 fold change) between AEP and 105 strains in their TE content, as a function of read count. **(d)** Representative locus with CR1 expression measured by Iso-Seq. Grey lines indicating Iso-Seq expression strength. **(e)** Tissue-specific expression of TEs in AEP lineage using both cell type resolved RNA-seq data^12^ and scRNA-seq^14^. **(f)** Cell-type composition analysis of specific TE expression. In the left upper panel, stem cell populations are colored in black and other differentiated cells colored in grey. The rest of panels show representative plots of A-TE subfamily expressions.

Next, we analyzed the expression profiles of TEs using the publicly-available single-cell RNA-seq dataset from adult polyps of the AEP strain^14^. We quantified the expression levels of TEs belonging to the 868 subfamilies that had been confirmed to be expressed by Iso-Seq in the whole polyp (Fig. 3a–d). A significant correlation was observed between the expression levels of TEs in Iso-Seq and scRNA-seq (R = 0.75, p < 2.0e-15; Fig. 3e). We investigated TEs that showed significantly higher expression levels in one or more of the six cell populations—comprising interstitial stem cells, cells differentiated from interstitial stem cells, ectodermal stem cells, cells differentiated from ectodermal stem cells, endodermal stem cells, and cells differentiated from endodermal stem cells—compared to the rest of the cell populations (Fig. 3f, left upper panel). At the family level, six types of TEs (TcMar-Tc1, CMC-Chapaev-3, PIF-Harbinger, hAT-Charlie, hAT-hAT5, and MULE-NOF) showed specific expression in one or more cell populations (Supplementary Data 5). All A-TEs were expressed at least in i-cells which is consistent with the fact that i-cells can give rise to germ line cells^12^, whose genomes are inherited by all three stem cell lineages through sexual reproduction (Supplementary Fig. 5b). At the subfamily level, 44 types of TEs showed cell-type-specific expression profiles, and at the loci-level analysis, 203 types of TEs exhibited cell-type-specific expression profiles (Supplementary Fig. 6, Supplementary Data 6,7). The expression profiles measured by scRNA-seq were also supported by FACS-sorted bulk RNA-seq datasets^12^ (Supplementary Fig. 6, Methods).

In summary, these data provide a comprehensive bulk, tissue, and cell-type resolved cross-validation analysis of Hydra TE expression dynamics, revealing distinct and cell-type specific patterns of TE expression and the high expression activity of the A-TE families.

### Identification of stem cell-specific transposon insertion sites using long-read sequencing

Hydra, primarily reproducing asexually through budding, can also resort to sexual reproduction under stress conditions such as low temperatures or starvation. In laboratory conditions, transgenic hydra strains are propagated asexually, with offspring inheriting genomes from three parental stem cell types. Over time, these strains may accumulate unique transposon insertions in each stem cell type’s genome. Utilizing this aspect of hydra biology, we investigated the genomic diversity caused by TE insertions in stem cells using the Cnnos1::GFP hydra transgenic line, which highlights interstitial stem cells with GFP expression, driven by the Nanos promoter (Fig. 4a)^26^. A clonal population propagated through budding from a single polyp for more than 10 years was subjected to FACS to isolate GFP-positive (i-cells) and GFP-negative cells (non-i-cells), followed by Nanopore PromethION long-read sequencing for each group, yielding 15.1 Gb and 16.3 Gb of long reads for i-cells and non-i-cells, respectively (Fig. 4b, Supplementary Fig. 7). These reads were aligned to the AEP genome sequence for variant calling with filtering (Supplementary Fig. 7, Methods).

**Fig. 4.**
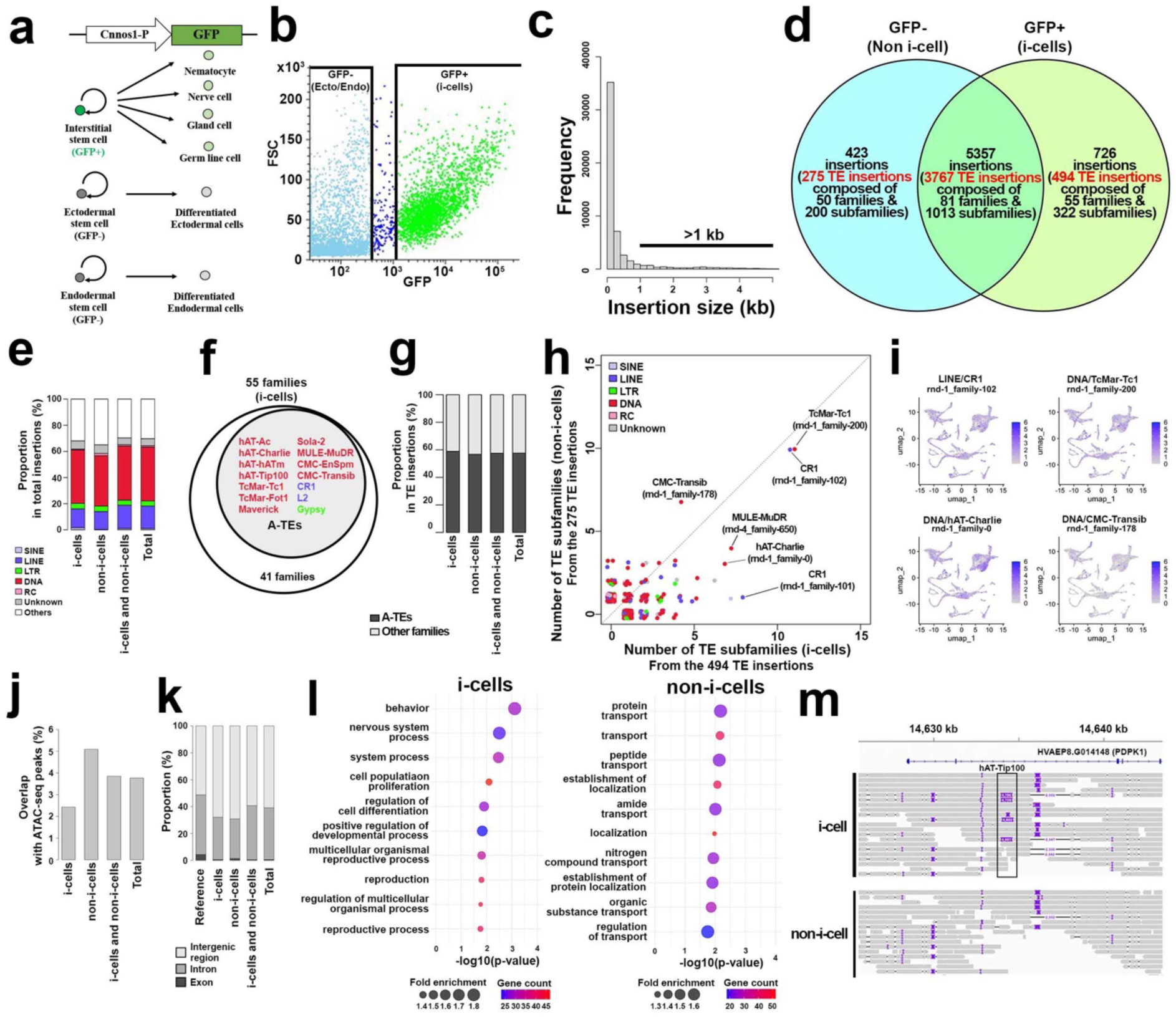
TE insertion dynamics of stem cells in Hydra. **(a)** Cnnos1-GFP transgenic line labelling i-cell stem cells used for this analysis. **(b)** FACS-sorting of the i-cell population and non-i-cell population. **(c)** Size distribution of insertions. **(d)** Number of cell type specific and shared insertions. Some of the insertions were further annotated as TEs. **(e)** Major types of TEs contributing to the insertions. **(f)** TE families which comprise the insertions of the i-cell population. The core 14 TE families are also shown. **(g)** Contribution of the A-TEs to the TE insertions. **(h)** Counts of TE subfamilies in the 494 TE insertions (x-axis) and 275 TE insertions (y-axis). **(i)** Expression profile of TEs which showed genomic insertions in the stem-cell lineage specific whole-genome sequencing. **(j)** Proportions of insertions overlapping open chromatin regions. **(k)** Distribution of insertions identified in intergenic regions, introns, and exons. **(l)** GO-term enrichment analysis of closest genes to the insertions detected in the i-cell as well as ecto/endo stem cell lineage genomes. **(m)** i-cell specific hAT-Tip100 insertion in the intron of 3-phosphoinositide-dependent protein kinase 1 (PDPK1) involved in cell proliferation^27,28^.

We identified 54,857 genomic insertions across both populations, with 6,506 larger than 1kb, indicative of active TE activity (Fig. 4c). Notably, 726 insertions were unique to i-cells, 423 to non-i-cells, and 5,357 were shared (Fig. 4d). TEs constituted a major part (69.7%) of these insertions, with significant representation from the A-TEs (Fig. 2e, 4e–g). Among the TE insertions, 68.0% (494 insertions) of the insertions were in the i-cell population, 65.0% (275 insertions) in the non-i-cell population, and 70.3% (3,767 insertions) shared by both populations, respectively (Fig. 4d). The 5,357 insertions including the 3,767 TE insertions which observed both in i-cells population and non-i-cell population can be regarded as reflecting the genetic background difference between the transgenic line and the AEP line used for the reference genome. Among these TE insertions, DNA elements were the most numerous, followed by LINE elements, and then LTR elements (Fig. 4e). These TE insertions were composed of 55 TE families, 50 TE families, and 81 TE families for the i-cell population, non-i-cell population, shared populations, respectively (Fig. 4d). Of note, all of the population included the A-TEs (Fig. 4f). At the subfamily level, the insertions in the i-cell population were impacted by 322 TE subfamilies; the non-i-cell population insertions by 200 TE subfamilies; and the shared insertions by 1,013 TE subfamilies (Fig. 4d). Of the TE insertions identified, 58.9% in the i-cell population, 56.7% in the non-i-cell population, and 57.5% in those found in both the i-cell and non-i-cell populations were attributed to the A-TEs suggesting the high activities of the A-TEs in each of the stem cell lineage (Fig. 4g). In both the i-cell and non-i-cell populations, the CR1 (rnd-1_family-102) and TcMar-Tc1 (rnd-1_family-200) subfamilies were the most prevalent TE subfamily insertions (Fig. 4h). In the scRNA-seq datasets of the hydra polyps^14^, the TEs showed relatively broad expression suggesting underlying potential contributions of chromatin states to the TE insertion preference (Fig. 4i). Upon examining the overlap between TE-insertions and publicly available ATAC-seq data^19^, we found that 2.5% of the TE insertions in the i-cell population, 5.1% in the non-i-cell population, and 3.8% of those identified in both populations overlapped with ATAC-seq peaks (Fig. 4j).

As TE insertions can drive variation and impact gene regulation, we examined the positional relationship between the TE insertions and protein coding loci. Insertions tended to preferentially accumulate in the intergenic regions (Fig. 4k). Analysis of the genes near the TE insertion loci specific to the i-cell population revealed 20 statistically significantly enriched gene ontology (GO) terms such as cell proliferation (left panel in Fig. 4l). This enrichment was highlighted by the i-cell specific A-TE hAT-Tip100 insertion in the intron of 3-phosphoinositide-dependent protein kinase 1 (PDPK1)^27,28^ (Fig. 4l,m). Similarly, 22 gene groups associated with TE insertions specific to non-i-cells were identified, with GO terms related to substance transport such as protein transport and peptide transport being significantly enriched (right panel in Fig. 4l).

Similarly, we performed stem-cell lineage specific whole genome sequencing on the ecto-GFP/endo-RFP line^29^, which expresses GFP in ectodermal epithelial stem cell lineage and RFP in endodermal epithelial stem cell lineage, and has been propagated by budding since 2011 (Supplementary Fig. 8a,b). We identified 409 TE insertions (54 families and 286 subfamilies), 523 TE insertions (61 families and 382 subfamilies), and 464 TE insertions (59 families and 345 subfamilies) specific to the ectodermal stem cell lineage cell population, endodermal stem cell lineage cell population, and non-epithelial cell population, respectively (Supplementary Fig. 8c,d). Similar to the TE insertion patterns in the Cnnos1::GFP line (Fig. 4f–h), these insertions were predominantly from the A-TEs (Supplementary Fig. 8e–g). However, they were associated with genes belonging to different GO categories, such as cilium organization in ectoderm, chromosome organization in the endoderm, and monoatomic ion transport in the non-epithelial cells (Supplementary Fig. 8h,i), suggesting different targeting of A-TE in distinct stem cell lineage genomes.

In summary, our lineage-specific whole-genome sequencing of stem cells has unveiled specific loci for TE insertions associated with distinct gene ontologies and predominantly influenced by the A-TEs. In addition, the contribution of the A-TEs to the general transgenic line differences suggests their accumulation in the germline.

### Deep evolutionary conservation of the A-TE families

Finally, we investigated whether A-TEs are present across a wide clade in the genomes of animals and their outgroups to test the phylogenetic scope of their putative activity beyond hydra. With a custom deep homology search (Methods), we analyzed TE homology and retention in 17 metazoans, three unicellular holozoans (*Creolimax fragrantissima*, *Capsaspora owczarzaki*, *Salpingoeca rosetta*), two fungi (*Candida auris*, *Saccharomyces cerevisiae*), one amoebozoa (*Dictyostelium discoideum*), and two plants (*Chlamydomonas reinhardtii*, *Arabidopsis thaliana*) (Fig. 5 and Supplementary Fig. 9). A total of 149 TE families were detected in at least one species analyzed (Fig. 5 and Supplementary Fig. 9). We discovered a cluster of several TE families with overrepresentation across the species analyzed (cluster #1 in Fig. 5 and Supplementary Fig. 9). We find that at least 64.3% (nine families) of these families are members of A-TEs (arrows in cluster #1 in Fig. 5 and Supplementary Fig. 9). The pan-eukaryotic distribution of these TE families suggests a very ancient conservation and roles of these A-TEs in shaping chromosomes that might predate the current metazoan-unicellular homologies^2^. We term these families eukaryotic Core-TEs. Besides the eukaryotic Core-TEs, many TEs were widely observed across metazoan (Supplementary Fig. 9), among several clusters we find that the other part of A-TEs are clustered within one of several pan-metazoan clusters (cluster #2 in Fig. 5 and Supplementary Fig. 9). While this cluster has a particularly high activity in the genus *Hydra*, these TE families are also abundant in other metazoan species and may thus represent an active metazoan Core-TE set.

**Fig. 5.**
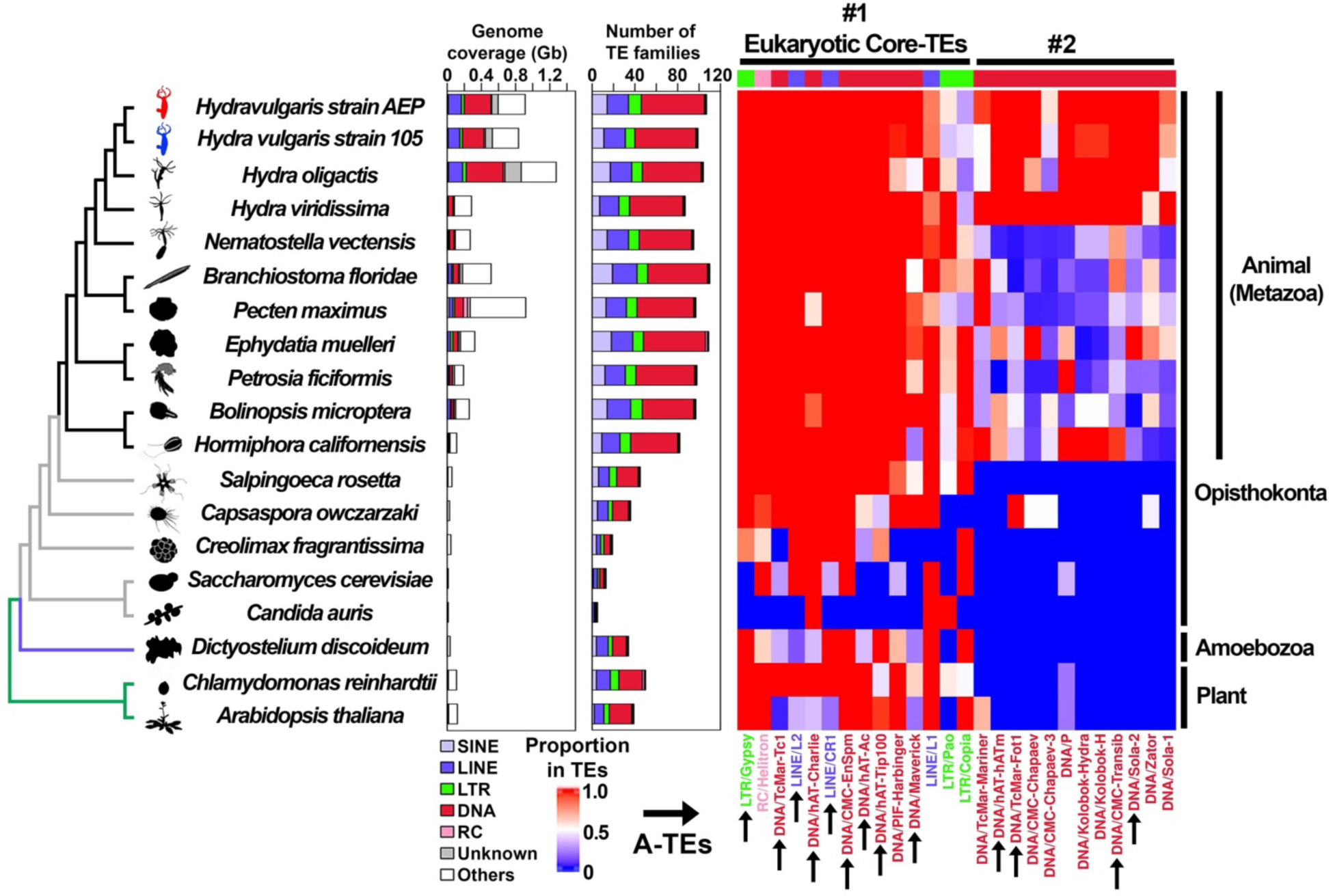
Deep TE homology and retention of genomically active TEs across eukaryotes. Compositions of selected TE clusters in eleven metazoan genomes and those of eight outgroup species. The complete results are shown in Supplementary Fig. 9. The species cladogram is based on the previous research^2,68–71^, and the edges of metazoans are colored in black and those of the outgroup species are colored in grey. Number of detected TE families are shown in a bar plot. TE families are expanded in metazoan species compared to the outgroup species. The heatmap shows the extent to which the genome coverage of each TE family constitutes the overall genome coverage of TEs in each species. The dendrogram on top of the heatmap represents the result of hierarchical clustering (Supplementary Fig. 9). The names of the TE families belonging to the clusters highlighted in #1 (eukaryotic core-TEs) and #2 (active in *Hydra*) are shown at the bottom. The 14 TEs of the A-TEs are indicated by arrows.

We find evidence that Core-TEs, including the eukaryotic core TEs, are continuously shaping animal chromosomes. We profiled the reported genome expansion in the zebrafish genome^30^ and found that the eukaryotic core TE cluster members are heavily involved in the *Danio rerio* genome expansion contributing to 30.0% of the genome. We find that the metazoan core cluster #2 is present but not as prominently active in the zebrafish genome as in the brown hydra genomes. Similarly, nine out of 14 core eukaryotic TEs have contributed to the ancient coleoid cephalopod genome expansion and evolution^31^. However, we also find other TE clusters that are expanded in species-specific manners, that are not covered by A-TEs and may thus constitute divergent lineage-specific active TE core sets that were not captured in our hydra-centric analysis.

Taken together, our data suggest that A-TEs are forming evolutionary conserved core sets of self-propagating elements on ancient chromosomes of eukaryotes and metazoans. The same core TE set has been employed multiple times in evolution contributing to independent genome expansions. Beyond the highly conserved eukaryotic core set, our data also suggest the presence of several other metazoan core TE sets (Supplementary Fig. 9), including one that contains the remainder of hydra A-TEs.

## Discussion

How TEs have shaped and continue to affect genomes of animals remains an understudied question due to the large variance in genomic representation of these elements, and, more crucially, the lack of model systems to profile their activity and function. To address this question, we utilized the cnidarian hydra with its distinct stem cell lineages that yield, under asexual reproduction, three independently evolving genomes within the same individual. With transgenic access to these genomes and cell type transcriptomics, we were able to reveal the putatively active TEs (A-TEs), the key TE components of these genomes and cell types, and to assess their expression and genomic insertion dynamics.

In this study, we provide direct evidence that the stem cells of hydra have diversified genomes due to specific sets of A-TEs. We show that on top of the TE family contributing to the genome-size expansion in the brown hydras such as the retroelement CR1, several other key and predominantly DNA elements are active in individual stem cell lineages. We show evidence for thousands of full-length TE transcripts, including evidence for trans-spliced leader and poly-A addition. Among these, over a dozen TE families are actively shaping genomic landscapes in interstitial, ectodermal, and endodermal stem cell lineages. Through inserting into sites in and around genes involved in various cellular functions these insertions are likely to contribute to generating variation in expression of these genes. For example, the observation of TE insertions near genes involved in cell proliferation may suggest a race between TEs to make their host cell outcompete others. This mechanism may be of crucial selective advantage for an asexually reproducing animal such as hydra.

TEs are prevalent in eukaryotic genomes^32^. Several superfamilies, such as Tc1/mariner and hAT,are known to be present across eukaryotes^32^. However, it was not well understood whether homologous TEs can be identified at the family level. It was also unclear if various TE families are randomly active in eukaryotic lineages, or if a limited number of TE families are widely conserved and active in eukaryotes. Detailed analysis at the family level has been conducted for specific metazoan clades such as mammals^33^, but not yet broadly in eukaryotes because of high substitution rates and fragmentations of inactivated TEs^34^. In this study, using a custom sensitive approach and a broad eukaryotic species sampling, we reveal that all of A-TEs are evolutionarily highly conserved either at the eukaryotic or metazoan levels, making them “core” evolutionarily preserved and active TEs of these genomes.

At the metazoan level, beyond the core set dominated by the A-TEs identified in this study, the evolutionary comparison also hints at the existence of a few other “core” metazoan sets. While the A-TE-enriched set is also the one showing highest contribution to the brown hydra genome expansion, its homology is widely conserved across metazoans. Similar “core” sets from this widely conserved metazoan pool of TEs may be employed in independent species-specific genome expansions. This suggests the possibility that animal chromosomes evolve through activation of particular core TE sets, while keeping the ancient chromosomal homologies stable. While the mechanistic insights into the evolutionary genome dynamics remains to be fully elucidated, the evolutionary conserved PIWI-piRNA pathway-medicated TE repression might be coupled with it^35^. Together, these findings open up an exciting opportunity to test how activations of these “core” TE sets are contributing to divergent macro-evolutionary trajectories of animal genomes.

## Methods

### Animal husbandry

Hydra strains were kept in individual containers filled with Hydra medium at 16 ℃. Hydras were maintained under a continuous cycle of 14 hours of light and 10 hours of darkness. Hydras received regular feedings of newly hatched brine shrimp. The Cnnos1::GFP strain and the ecto-GFP/endo-RFP strain were from previous studies^26,29^.

### Generation of the AEPx105 strain

Males of the AEP Hym176B promoter::GFP line from Toshitaka Fujisawa were crossed with NE07a females. One of the progenies from this cross carried the transgene; this animal was used to establish a line. This line was heterozygous for the transgene. Females from this line were crossed with males from the Landing Site 6.1 line, which is an AEP transgenic line homozygous for the actin promoter::DsRed2 transgene and is six generations inbred. From this cross, lines were established that carried both transgenes (heterozygous for GFP and heterozygous for DsRed2). Males from these lines were crossed to females from the SS1 AEP line (an AEP F1 line that is “supersexy”). Subsequent studies using progeny from these crosses showed that the transgenes were on the same chromosome and that in one of these progenies a crossover had occurred that linked the two transgenes on that chromosome. This line was 7/8 AEP and 1/8 NE07a, and it was heterozygous for the chromosome containing the linked transgenes. Males from this line were crossed with 105 females to generate the AEPx105 hybrid strain. The AEPx105 strain that was obtained did not carry the transgenes.

### Nanopore Sequencing

Extraction of high molecular weight DNA from a snap-frozen sample of hydra polyps of the AEPx105 strain and Nanopore sequencing on an Oxford Nanopore Technologies PromethION were carried out by the DNA Technologies and Expression Analysis Cores at the UC Davis Genome Center, which is supported by NIH Shared Instrumentation Grant 1S10OD010786-01. Following the initial run, the flow cell was given a nuclease flush and reloaded for a second round of data collection. The two datasets were combined for subsequent use.

### Omni-C Sequencing

A snap-frozen sample of hydra polyps of the AEPx105 strain was provided to Cantata Bio (Scotts Valley, California) who produced and sequenced an Omni-C library.

### Iso-Seq Sequencing

RNA was extracted from hydra polyps of the AEPx105 strain using the method of Chomczynski and Sacch^36^. Following resuspension in DEPC-treated water, the RNA was precipitated with 2.5 M LiCl overnight at 4°C. The RNA was pelleted, washed twice with 70% ethanol, resuspended in DEPC-treated water and stored at -80°C. Synthesis of cDNA, preparation of the SMRTbell template, and sequencing (two SMRT cells) were done at the Genomics Research and Technology Hub (GRT Hub) at UC Irvine on a Pacific Bioscience Sequel II sequencer.

### Generation of haplotype-resolved genome assembly of the AEPx105 F1 hybrid

The heterozygosity of the AEPx105 F1 hybrid genome was assessed from the Illumina short reads of the gnome^1^ using Jellyfish v2.3.1 ^37^, a k-mer counting software, with a k-mer size of 21. The k-mer spectrum was analyzed using GenomeScope^38^. The resulting k-mer profile displayed a single peak, suggesting that the AEP haplotype and the 105 haplotype were well diversified and resolving each haplotype genome is feasible. In the previous study, total 4,420 contig sequences with the total size of 1.75 Gb were previously assembled from 23.6 Gb PacBio HiFi reads of the AEPx105 genome^1^. The chromosome-level scaffolds were constructed with the combination of the 4,420 contig sequences and the newly generated 26.2 Gb DNase I Hi-C sequencing reads. The read quality of the Hi-C reads were evaluated using FastQC v0.12.1. The Hi-C reads were mapped to the contig sequences using Juicer v1.6^39^ with default parameters. Subsequently, Hi-C scaffolding was performed using the 3D-DNA pipeline v180922^40^ with parameters the default parameters followed by curation of resulting scaffolds using Juicebox Assembly Tools v1.11.08^41^. This Hi-C scaffolding step produced 30 chromosomal scale scaffolds which are corresponding to both of 15 haplotype chromosome sets. These scaffolds comprise 97.1% (1.70 Gb) of the initial contigs. By comparing with previously reported genome assemblies of the AEP strain^19^ and the 105 strain^1^, we confirmed that 15 of the scaffolds correspond to the paternal AEP haplotype genome, while the remaining 15 are correspond to the maternal 105 haplotype genome. The AEP haplotype genome sequences had 1,523 gaps, while the 105 haplotype genome sequences had 2,300 gaps. Subsequently, all these gaps were filled with the newly generated 48 Gb of raw Oxford Nanopore reads. We identified telomere reads from the previous HiFi reads and newly generated Nanopore reads for this study and extended the terminal sequences of chromosomes. Resulting scaffolds were polished with previously obtained 35.6 Gb Illumina short reads of the AEPx105 F1 hybrid^1^. In the final genome assembly, each haplotype was gapless and featured telomere-subtelomere sequences at both ends of all 15 chromosomes. The AEP haplotype genome was 912 Mb in size, while the 105 haplotype genome was 834 Gb in size. Compared to previously reported data, the AEP haplotype exhibited an increase of 10.9 Mb, while the 105 haplotype showed an increase of 16.2 Mb. Completeness of the genome assemblies examined using BUSCO v5.4.6 and the metazoa_odb10 datasets^42^. The BUSCO scores on genome sequences for both haplotype genomes were 91.7%, an improvement over the previously reported AEP genome assembly and 105 genome assembly scores of 91.5% and 91.4%^1,19^. Based on these findings, the genome assembly appeared closer to a complete genome assembly. We have named this genome assembly HydraT2T.

### Analysis of repetitive elements of the reference genome assembly

Sine the majority of TEs are inactivated and degenerated due to accumulation of mutations such as base substitutions, insertions, and deletions, TE annotation in a pan-metazoan scale is challenging^34,43^. At the first step of TE annotation, generating custom repeat libraries using RepeatModeler v 2.0.5^44^ is a standard approach. A simple RepeatModeler run occasionally produces a library that contains lots of repetitive elements as ’Unknown’ elements. Analyzing the genome sequence with RepeatMasker v4.1.6 using this custom repeat library, which includes many ’Unknown’ sequences, inevitably results in a large number of repetitive elements being defined as ’Unknown.’ Since the process of RepeatModeler^44^ generating consensus sequences for repetitive elements does not refer to the TE database such as Dfam^45^, RepeatMasker using RepeatModeler custom library still detects the proper proportions of repetitive elements in the genome sequence. Therefore, for the consensus sequences generated by RepeatModeler, we performed annotation using the nhmscan v3.4^46^, which uses a more sensitive hidden Markov model instead of rmblast, which RepeatClassifier uses. We searched for the best hit in the Dfam 3.8 database^45^ for each consensus sequence and assigned it as the TE annotation. With the annotated custom repeat library, RepeatMasker was performed on the genome sequences^47^.

### Genome annotation

To predict protein-coding genes in the genome assembly, we aligned Iso-Seq reads to the genome assemblies using the minimap2 v2.26-r1175^48^ with options -ax splice -uf -C5. We then performed genome annotation using BRAKER2 v 2.1.6^49^ on the repeat-masked genome assemblies masked by RepeatModeler^44^ and RepeatMasker^47^. The UTR sequences were further added to the gene prediction with the parameters of --addUTR=on –skipAllTraining.

### Synteny analysis

Reciprocal blast best hits were used to define orthologs between species. Ancestral linkage groups were identified and plotted using the R package macrosyntR v0.3.3 with default parameters^50^. Collinear synteny blocks were identified using MCScanX with default parameters^51^. Collinear synteny blocks were visualized using SynVisio^52^.

### Stem cell lineage specific whole genome sequencing

One hundred Hydra individuals with clonal propagation were carefully collected and transferred to a 2.0 ml tube containing dissociation medium^53^. The Hydra were then gently dissociated by incubating them in Pronase solution (0.1% Pronase in Hydra medium) for 90 min at 18 ℃. Throughout the incubation phase, gentle agitation and pipetting were performed to facilitate the disassociation procedure. After the incubation, the Pronase activity was neutralized by adding an equal volume of BSA-containing Hydra medium. The dissociated cells were subsequently washed three times with 1 ml dissociation medium. Finally, the dissociated cells were collected by centrifugation at 2000 rcf for 3 min at 4 ℃ and then resuspended in 1 ml dissociation medium. FACS were performed with BD FACSMelody™ Cell Sorter at the Max Perutz Labs BioOptics FACS Facility. The FACS sorted cells were suspended in a lysis buffer (pH 8.0) containing 10 mM Tris, 100 mM EDTA, and 0.5% SDS to lysis the cell and solubilize genomic DNA. Equal volume of phenol:chloroform:isoamyl alcohol (25:24:1, v/v/v) was added to the sample lysate, and the mixture was vortexed for 10 seconds to ensure proper emulsification. After mixing, the suspension was centrifuged at 12,000 rpm for 5 minutes at 4°C to facilitate phase separation. The upper aqueous phase contining genomic DNA, was carefully transferred to a new tube, avoiding the interphase. Equal volume of chloroform was added, and mixture was vortexed for 10 seconds. The suspension was centrifuged at 12,000 rpm for 5 minutes at 4°C. The upper aqueous phase containing genomic DNA, was carefully transferred to a new tube, and the same step was repeated once. The final aqueous phase was then treated with equal volume of 2.5 volumes of room temperature Isopropanol and 1/10 volume of 3 M sodium acetate (pH 5.5). The sample was mixed well. After incubation, the sample was centrifuged at 6,000 rcf for 5 minutes at 4°C. The supernatant was discarded, and the pellet was washed with 70% ethanol, air-dried three times, and finally resuspended in 100 μl TE buffer. The concentration and purity of the extracted genomic DNA were determined using a Nanodrop spectrophotometer. Library preparation and nanopore sequencing were performed at the Next Generation Sequencing core facility at the Vienna BioCenter.

### Nanopore read mapping to the genome sequence and variant calling for the stem-cell lineage whole genome sequencings

Qualities of the Nanopore reads were evaluated using NanoPlot^54^. Nanopore reads were aligned to the Hydra reference genome sequence^19^ using minimap2 v2.26-r1175 with options of -ax map-ont--secondary=no^48^. Variant calling for each individual sample was performed using Sniffles v2.02^55^ with the --minsupport 2 option which produces a SNF file for each sample. Individual variant calls were merged using Sniffles v2.02 with options of --combine-low-confidence 0.01 --combine-low-confidence-abs 1 --combine-null-min-coverage 3 which produce a combined VCF file. The genomic insertions where all samples have genotypes were kept using bcftool v1.19^56^. The insertions were further subset by bcftools v1.19 based on the combination of genotypes for each sample, for the downstream analyses. For insertion annotations, sequences of insertions longer than 1kb were extracted and searched against the RepeatModeler repeat library using NCBI BLASTN^57^. The TE annotation of an insertion was determined based on the consensus sequence of the custom repeat library that formed an alignment of 500 bp or more with the TE sequence.

### Functional gene analysis

For functional gene analysis, protein sequences were retrieved from the genome sequence and the genome annotations using GffRead v0.12.7^58^. The longest protein sequences were analyzed to identify homologous sequences within the NCBI-nr database, employing the NCBI BLASTN program with an e-value threshold of 1e-05. The longest protein sequence for each gene was functionally annotated using eggNOG-Mapper v2.1.9^59^. Genes proximal to TE insertion sites were identified using BEDTools v2.30.0^60^ with a subcommand of closest. We conducted Fisher’s exact test for the over-representation of GO terms within the group of genes proximal to TE insertion sites, utilizing the R library topGO v 2.54.0^61^. Results of the statistical tests were plotted using the R library ggplot2 v 3.5.0^62^.

### RNA-seq analysis

To determine whether each Iso-Seq read is derived from TEs, each read sequence from Iso-Seq was searched against the repeat library generated by RepeatModeler^44^ using NCBI BLASTN^57^ with default parameters. Iso-Seq reads that formed alignments of over 1 kb with the sequences in the repeat library were called as TE-derived sequences. Furthermore, Iso-Seq reads defined as TE-derived were mapped to the genome sequence using Minimap2 v2.26-r1175 with parameters of - ax map-hifi --secondary=no, and the mapped regions were identified. Additionally, to reveal the transcription of TEs at a single-cell resolution, sequencing reads from a publicly available scRNA-seq dataset of hydra^14^ were downloaded. From this dataset, only TE sequences belonging to the TE subfamilies confirmed to be expressed by Iso-Seq and that were over 1 kb in length were used. With these TE sequences, the expression levels of TEs in each cell were quantified using Alevin v1.2.1^63^. The resulting expression matrix was loaded using the CreateSeuratObject function of the R package Seurat v5^64^. Cell type annotation was based on the previous reports^14,19^. TEs showing cell type-specific expression were identified using the FindAllMarkers function of Seurat. Additionally, RNA-seq datasets for FACS-sorted cell populations^12^ were mapped to the genome sequence using HISAT2 v 2.2.1^65^. The read counts of each TE were evaluated with featureCounts^66^, and transcripts per million were calculated for each TE. The expression of TEs showing cell type-specific expression obtained from scRNA-seq was checked. Heatmaps were plotted using the R package ComplexHeatmap v 2.18.0^67^.

## Data availability

The sequencing reads are available in the NCBI database under BioProject accession number PRJNA1085048. The genome assemblies for the AEP and 105 genomes are available in the NCBI database under BioProject accession numbers PRJNA1085304 and PRJNA1085315, respectively.

## Code availability

The scripts for this manuscript are available at GitHub (https://github.com/ironman-tetsuo/Hydra_TE).

## Acknowledgments

The authors would like to express their gratitude to Yoshihiro Omori from Nagahama Institute of Bioscience and Technology, Japan, for his advice on genome extraction and to Juan Daniel Montenegro Cabrera of the University of Vienna, Austria, for his comments regarding repeat annotation of the Hydra genome. Our appreciation also extends to Nina Znidaric, Angela Caballero Alfonso, and Eduard Renfer of the University of Vienna for their dedication to animal care and technical assistance, as well as to Chiemi Nishimiya-Fujisawa from Heidelberg University, Germany, for her valuable advice on Hydra strains. We are thankful to Bert Hobmayer of the University of Innsbruck, Austria, for his insightful comments on this project. Whole genome sequencing was expertly performed by the Next Generation Sequencing core facility at the Vienna BioCenter. Special thanks to Kitti Dora Csalyi and Thomas Sauer at the Max Perutz Labs BioOptics FACS Facility for their valuable advice on the FACS experiments. Most of the computations were carried out on the Life Science Compute Cluster (LiSC) of the University of Vienna. R.E.S. thanks Dr. Ira Blitz (UC Irvine) for his assistance with RNA isolation for Iso-Seq sequencing. We thank all members of the Simakov lab, especially Darrin Schultz for valuable comments on the manuscript.

## Funding

This work was supported by the Austrian Science Fund FWF (I4353) to OS and EMT, D-A-CH (Ho1088-10-1) to TWH, Takeda Science Foundation to TK, the Mochida Memorial Foundation for Medical and Pharmaceutical Research to TK, the International Medical Research Foundation to TK, the Yamada Science Foundation to TK. O.S. was supported by an ERC-H2020-EURIP grant, grant number 945026. This work utilized resources of the UCI Genomics Research and Technology Hub (GRT Hub) parts of which are supported by NIH grants to the Comprehensive Cancer Center (P30CA-062203) and the UCI Skin Biology Resource Based Center (P30AR075047) at the University of California, Irvine, as well as to the GRT Hub for instrumentation (1S10OD010794-01and 1S10OD021718-01).

## Inclusion & Ethics

Every contributor to this research has met the authorship requirements and has been credited as an author, given their crucial role in conceptualizing and executing the study. The research encountered no significant limitations or prohibitions within the researchers’ environment.

## Conflicts of interest

The authors declare that they have no conflicts of interest.

---

**Supplementary Fig. 1.**
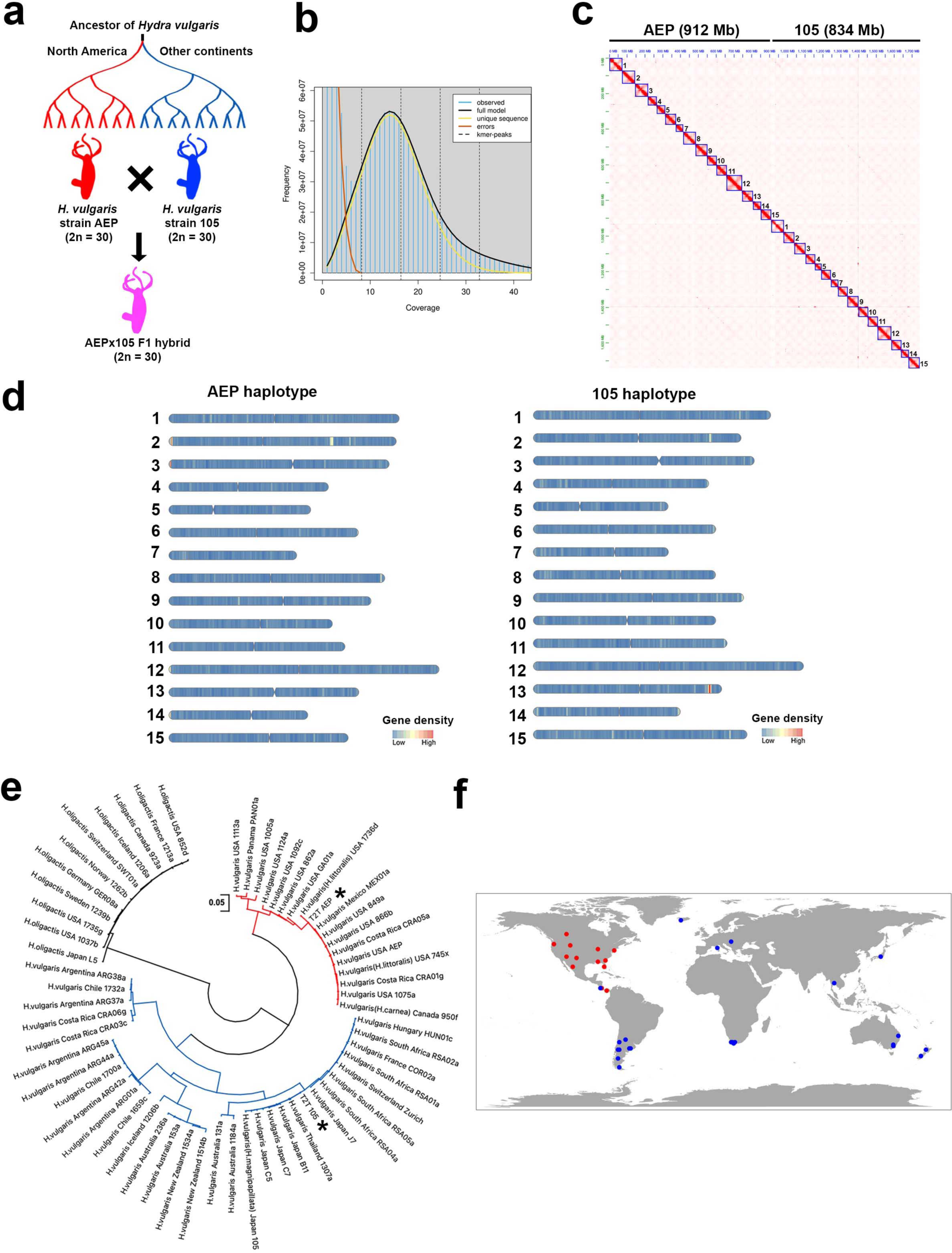
The 30 chromosomes of the AEPx105 F1 hybrid. **(a)** Generation of AEPx105 F1 hybrid by crossing the AEP strain and the 105 strain. **(b)** The k-mer frequency distribution of the AEPx105 F1 hybrid genome. The k-mer (k = 21) occurrence was analyzed in the raw short-reads of the AEPx105 F1 hybrid genome^1^. **(c)** Hi-C contact map of the 30 chromosomes of the AEPx105 F1 hybrid. In each chromosome, interaction signals at both ends of the chromosome are observed, suggesting a Rabl-like configuration, an arrangement of interphase chromosomes where centromeres and telomeres are located at opposing poles within the nucleus. **(d)** Chromosomes are colored according to gene density. The positions of the centromeres are indicated by notches. It is evident that each haplotype genome consists of metacentric or submetacentric chromosome. **(e)** Molecular phylogenetic positions of the AEP haplotype and the 105 haplotype. Maximum-likelihood tree based on sequences of the internal transcribed spacer (ITS) 1, the 5.8S ribosomal RNA gene, and ITS2 from the AEP haplotype, the 105 haplotype, and sequences from a public database. Samples of *Hydra oligactis* are colored black. Two major clades of *Hydra vulgaris* are colored red and blue. The ITS1-5.8S-ITS2 sequences of the AEP haplotype and 105 haplotype are indicated by asterisks. **(f)** Geographic distribution of the samples whose ITS1-5.8S-ITS2 sequences are publicly-available. Each dot represents individual sample, colored similarly to the Maximum-likelihood tree.

**Supplementary Fig. 2.**
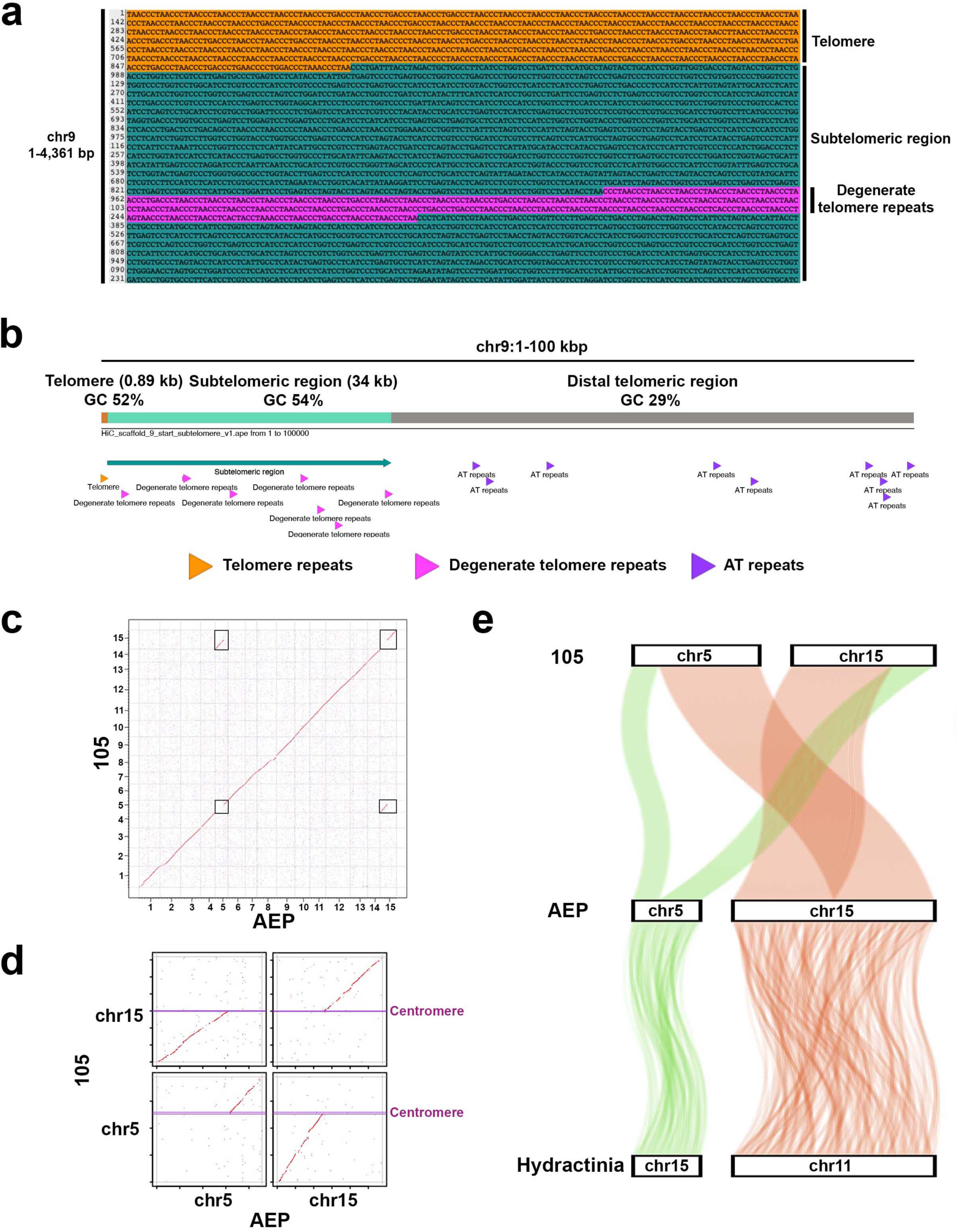
The structure of hydra telomeric, subtelomeric and centromeric regions. **(a)** Telomere and subtelomere sequences in the 1-4,361 bp region of the chromosome 9. **(b)** Motif map of the telomere, subtelomere, and distal telomeric regions in chromosome 9. **(c)** Genome alignment between the AEP chromosomes and the 105 chromosomes. Alignments of the chromosomes involved in the translocation is highlighted with rectangles. **(d)** Genome alignment between chromosome 5 and chromosome 15 from the AEP haplotype genome and those from the 105 haplotype. The start and end of the centromeric regions are indicated by horizontal lines. **(e)** Highlight of the Robertsonian translocation. Each curved line indicates positions of orthologs.

**Supplementary Fig. 3.**
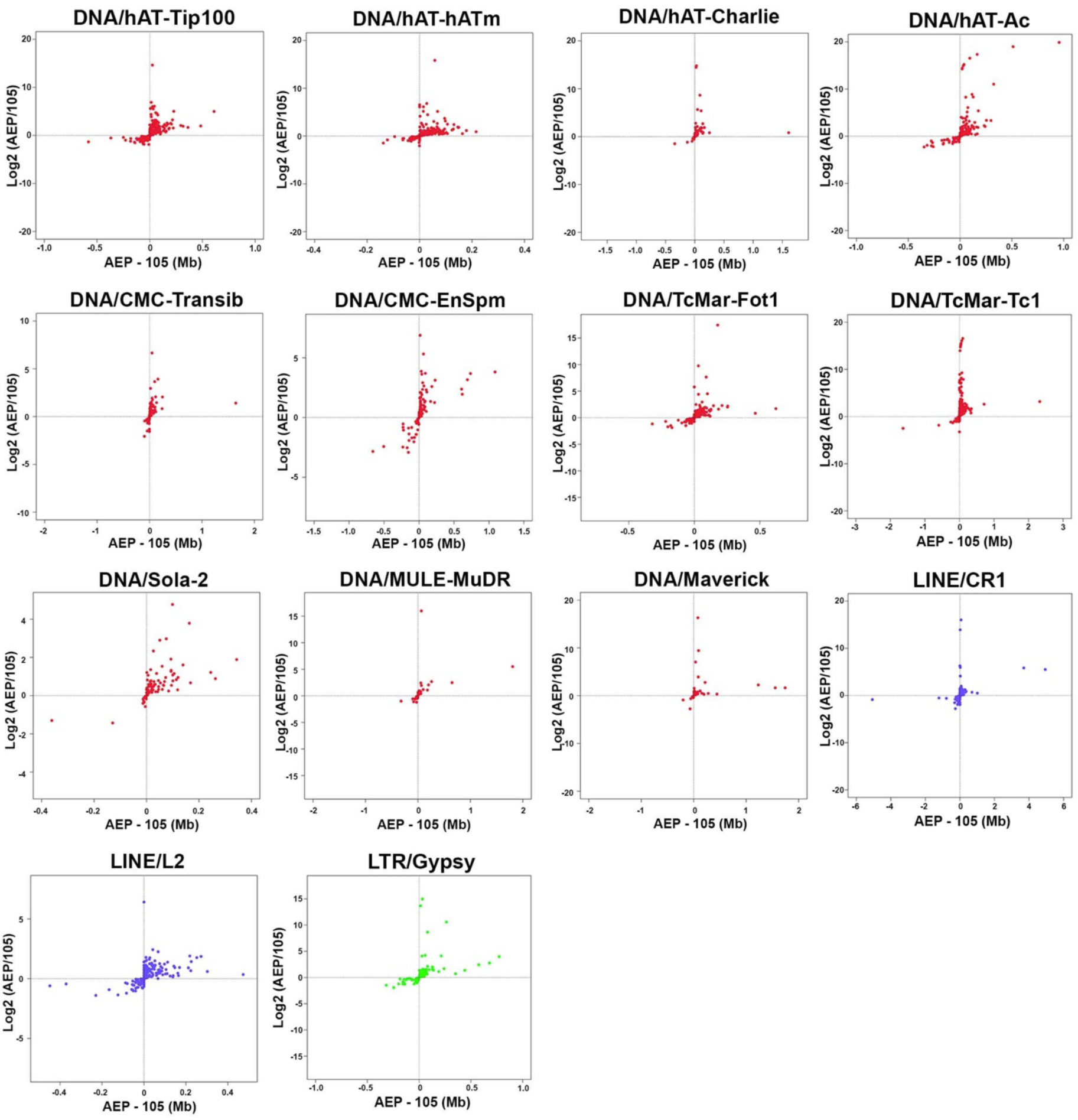
Divergence of the genome coverage of subfamilies of A-TEs between the AEP and 105 genomes. The x-axis represents the difference in genome coverage between each subfamily in the AEP genome and that of the 105 genome. The y-axis represents the ratio of genome coverage of each subfamily in the AEP genome to that of the 105 genome.

**Supplementary Fig. 4.**
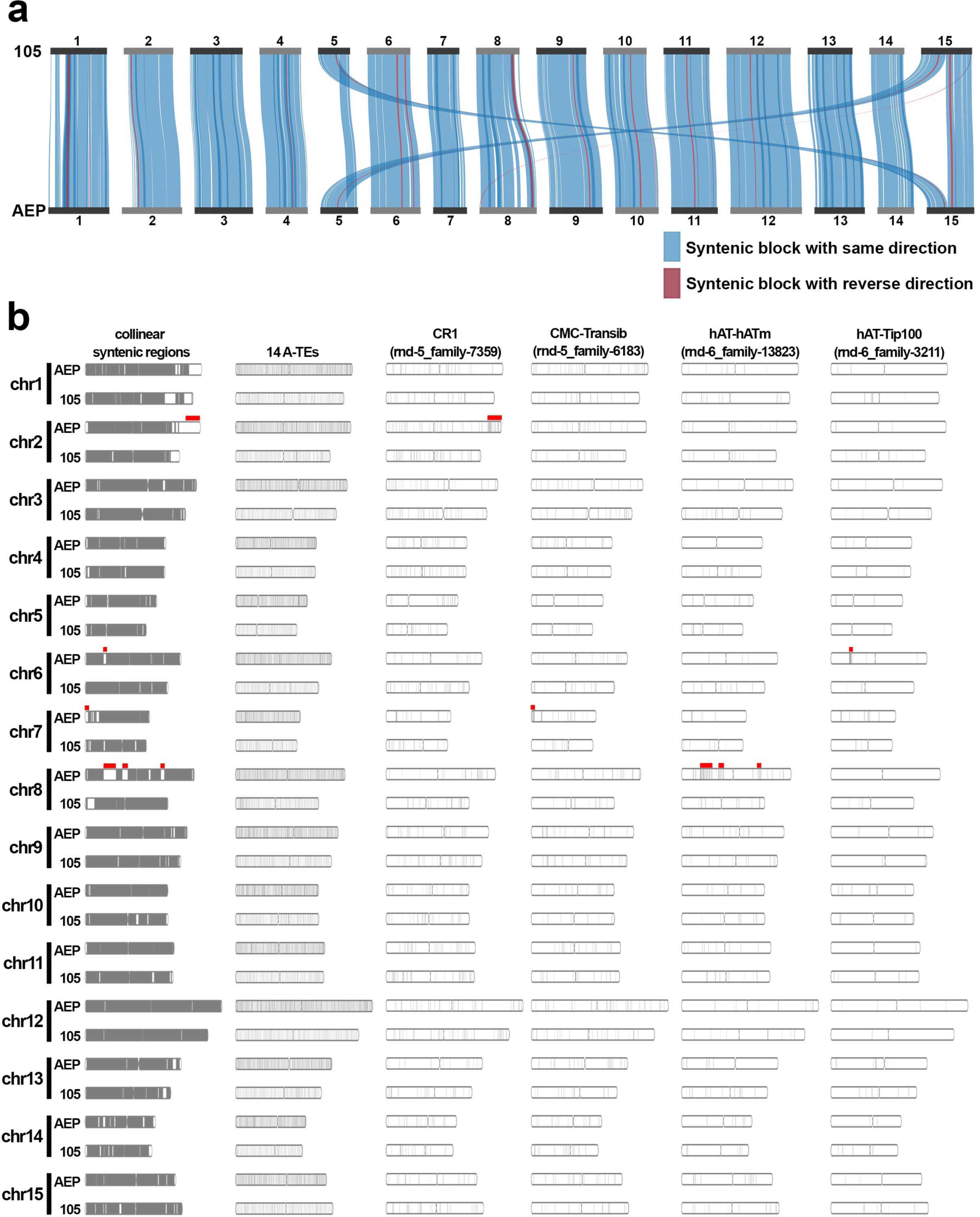
Divergence of TE subfamilies between the AEP and 105 genomes. **(a)** Collinear syntenic blocks between the 105 chromosomes and the AEP chromosomes. Collinear syntenic blocks are shown as ribbon diagrams. Syntenic blocks with same chromosomal orientations are colored in blue. Syntenic blocks with reverse orientations are colored in red. The length of the chromosomes is depicted as proportion to the assembly size of each chromosome. **(b)** Distributions of TE subfamilies across chromosomes. The leftmost panel shows the distribution of collinear syntenic regions in black. The second panel from the left represents the density plot for the 14 A-TEs. The remaining panels provide examples of the distributions of subfamilies of A-TEs. The red lines indicate genomic regions where collinear syntenic regions were not detected.

**Supplementary Fig. 5.**
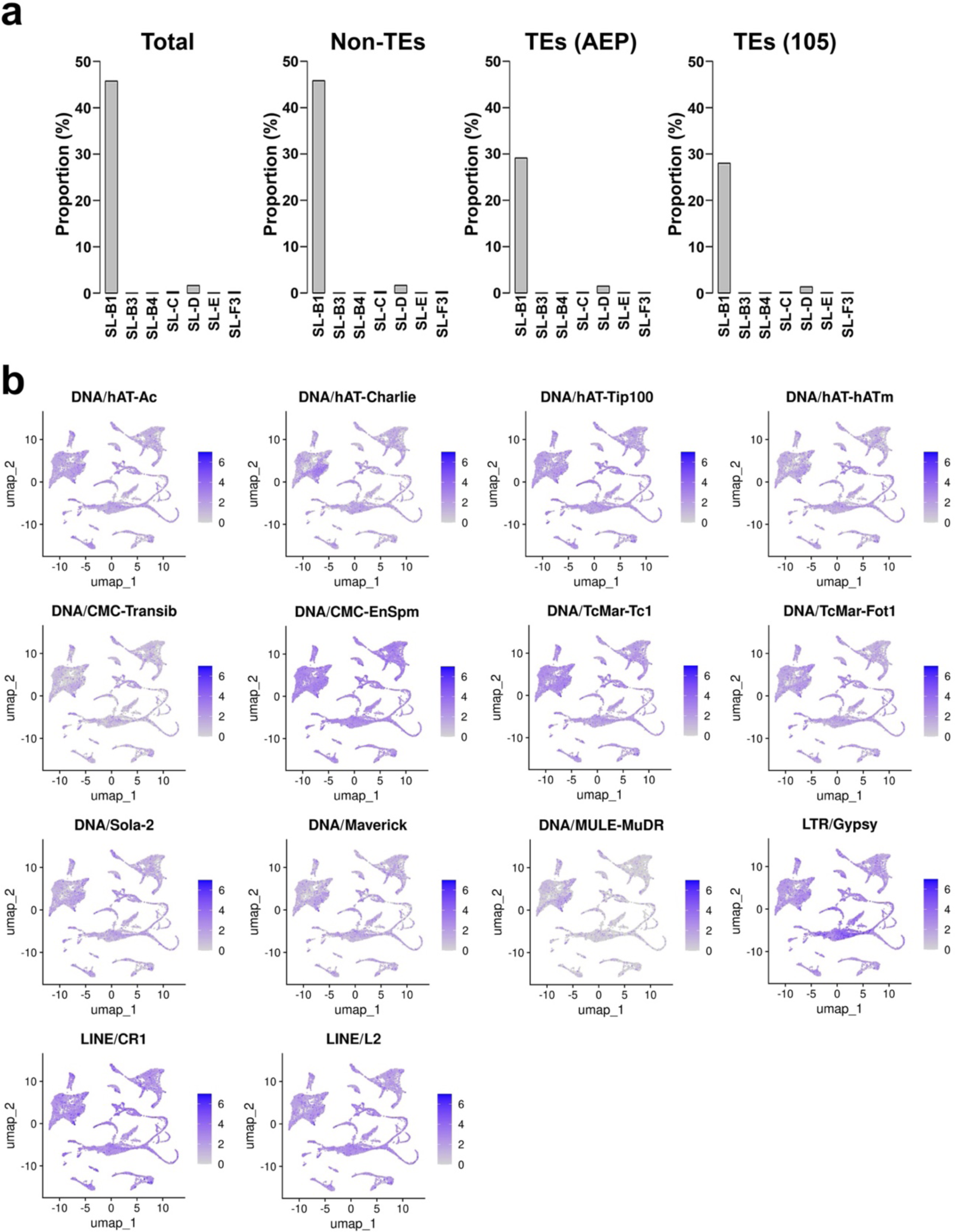
Expression profiles of TE families. **(a)** Proportions of transcripts with trans-spliced leader sequences revealed by Iso-Seq. The x-axes represent types of trans-spliced leader sequences^16^. The y-axes represent the proportions of transcripts with trans-spliced leader sequences. **(b)** Expression profiles of the A-TEs at the family level. The color scale indicating gene expression levels is the same in all panels, and the variations in color intensity are comparable across different panels.

**Supplementary Fig. 6.**
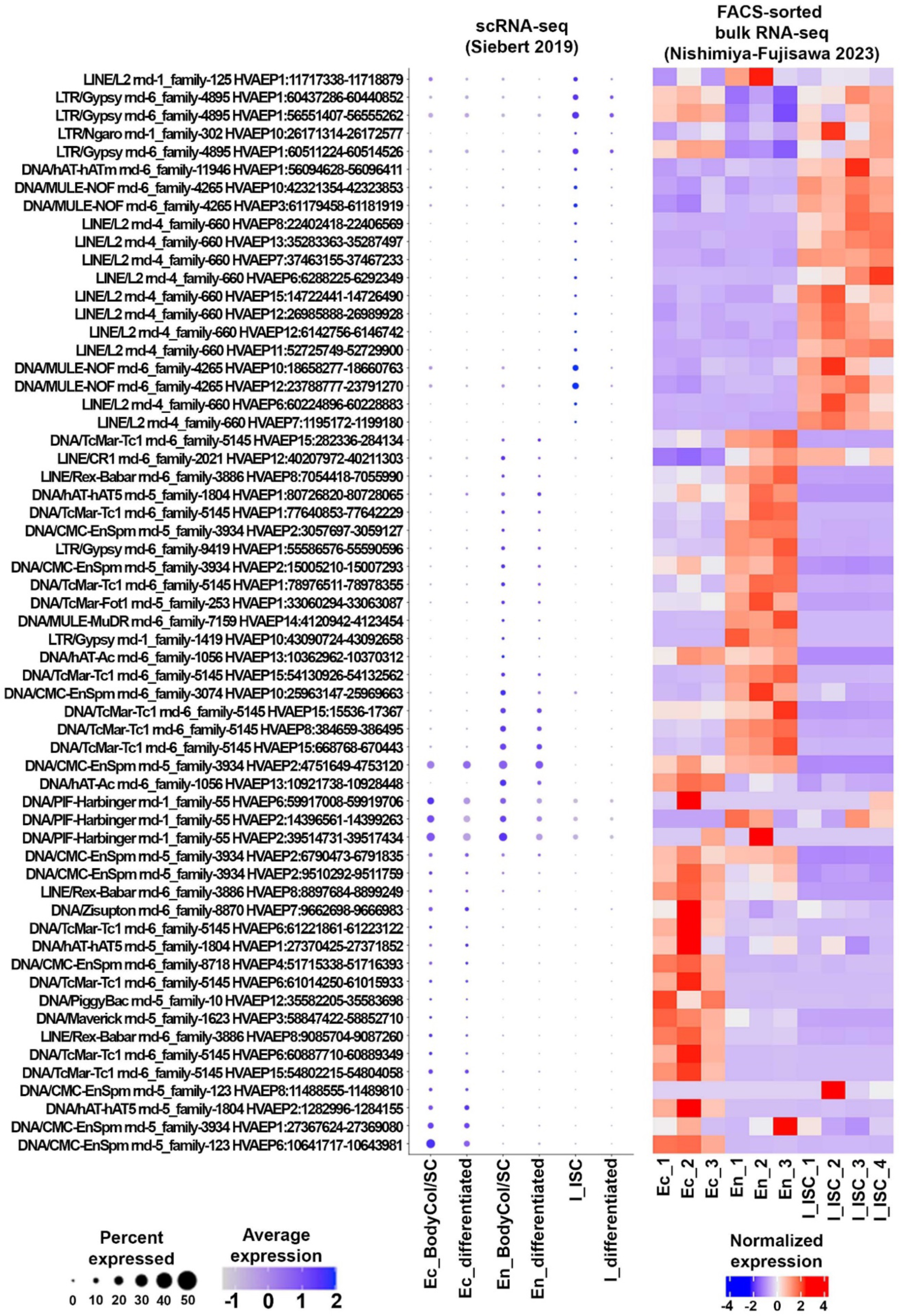
Expression profiles of TEs at locus level. Expression profiles of TEs at locus level were investigated using two different datasets^12,14^. In each group of ectodermal stem cells (Ec_BodyCol/SC), endodermal stem cells (En_BodyCol/SC), and i-cells (I_ISC), top20 TEs that were significantly more expressed are shown. Average expression levels (color intensity) and the proportion of cells expressing them (circle size) are shown. Additionally, as a validation, the expression levels of these TEs in a FACS-sorted bulk RNA-seq dataset are displayed on a heatmap to the right.

**Supplementary Fig. 7.**
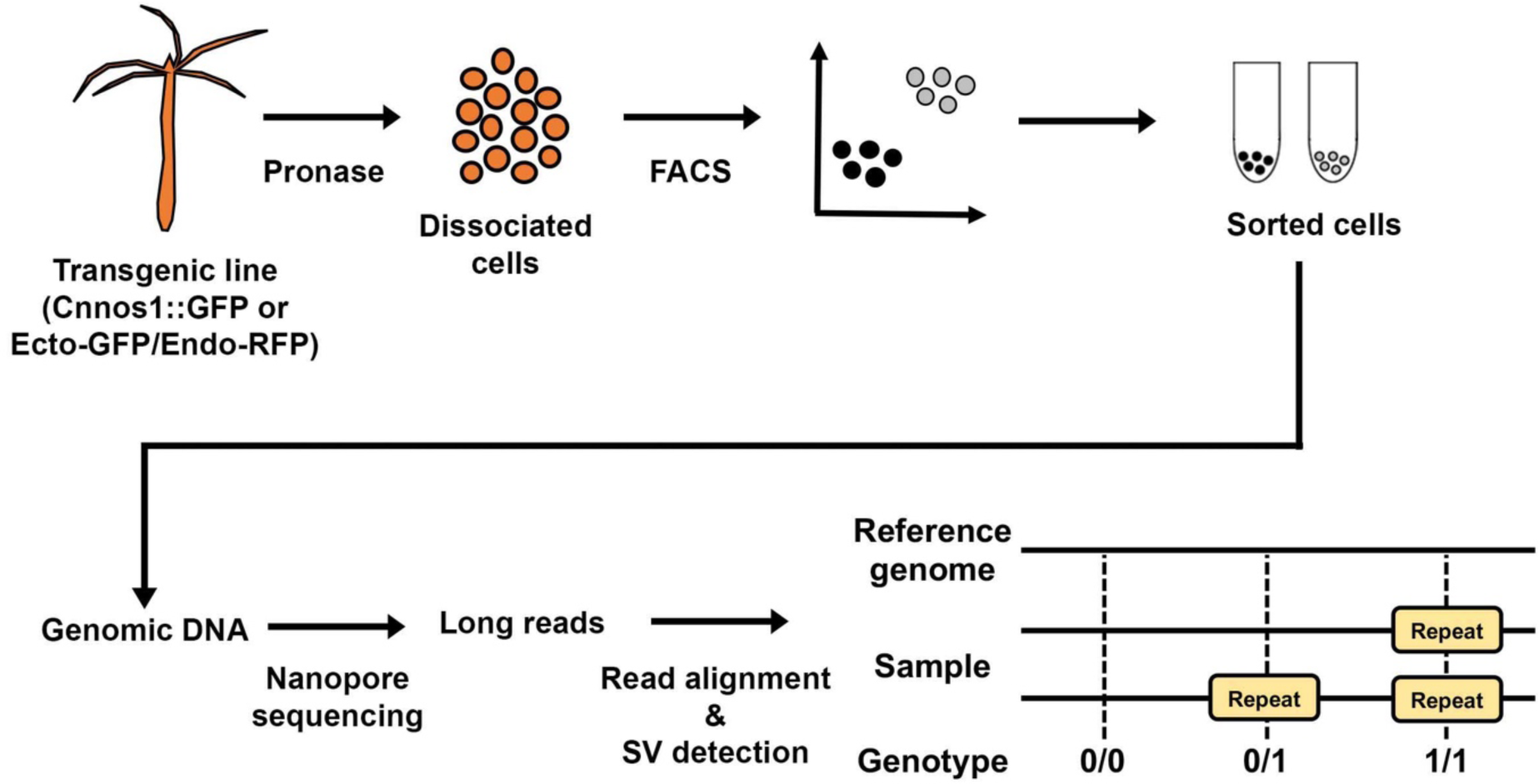
Workflow for stem cell-lineage specific whole genome sequencing. To elucidate the TE landscapes within the genomes of three stem cell lineages in hydra, transgenic lines expressing reporter genes (Cnnos1::GFP^26^ or Ecto-GFP/Endo-RFP^29^) were clonally propagated. The Hydra specimens were dissociated into single cells through Pronase treatment, and the stem cell lineages were sorted based on fluorescence intensities using FACS. Genomic DNA was then extracted from the sorted cells, and long reads were obtained via Nanopore sequencing. The reads were mapped to the AEP genome sequence to identify insertion sites. The analysis focused on insertions longer than 1kb, searching for insertions with sequences homologous to a custom repeat library generated by RepeatModeler from the genome sequence (Methods). This approach allowed for the determination of the extent to which each TE subfamily is inserted into the genome across different cell populations, facilitating the analysis of TE-induced stem cell genome divergence through comparisons across cell populations.

**Supplementary Fig. 8.**
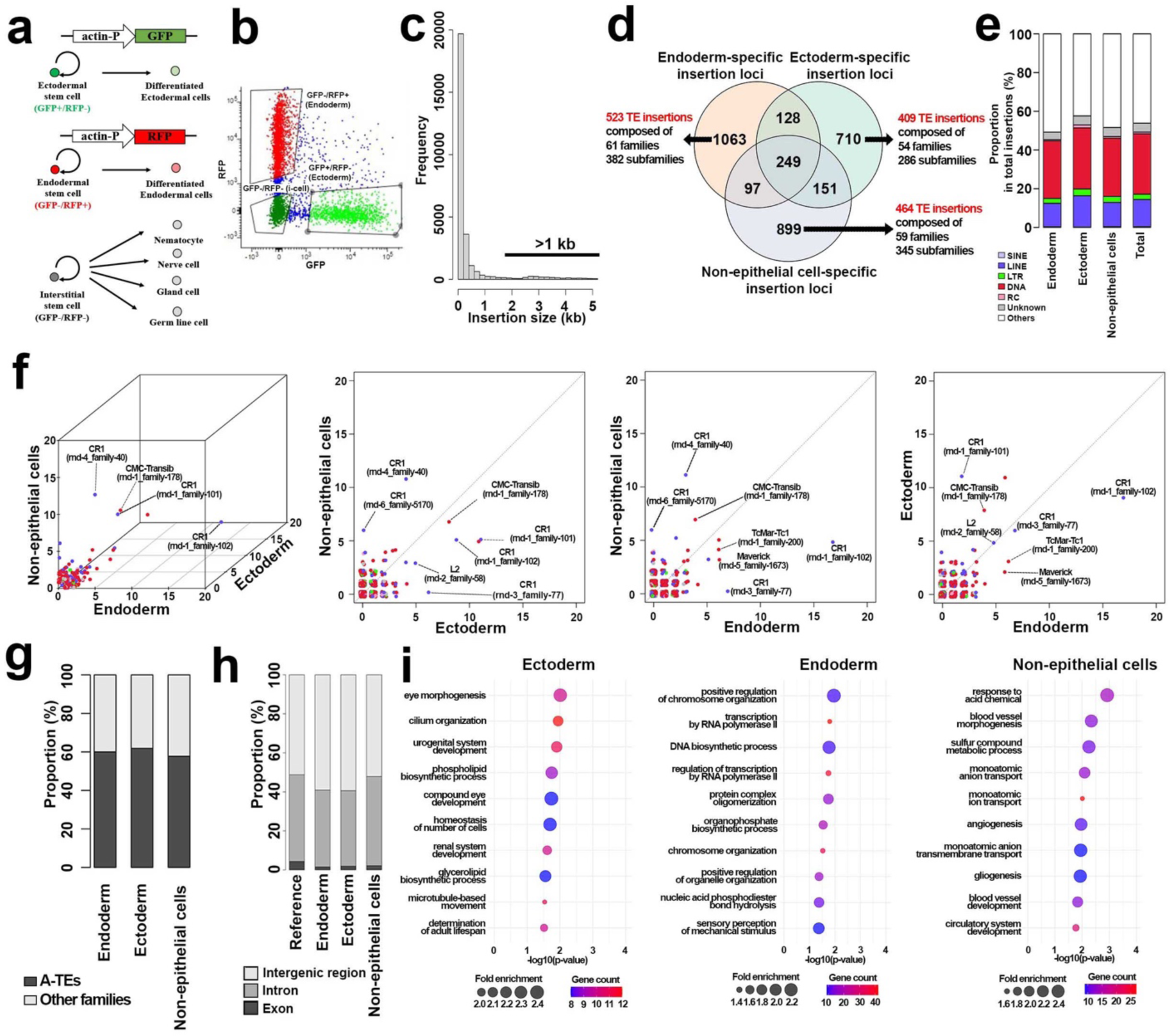
TE insertion dynamics of stem cells in ecto-GFP/endo-RFP hydra. **(a)** ecto-GFP/endo-RFP transgenic line labelling the epithelial cells. **(b)** FACS-sorting of the ecotodermal cell population, endodermal cell population and non-epithelial cell population. **(c)** Size distribution of insertions. **(d)** Number of cell type specific and shared insertions. **(e)** Major types of TEs contributing to the insertions. **(f)** Counts of TE subfamilies in the TE insertions at the specific loci for each population. **(g)** Contribution of the A-TEs to the insertions. **(h)** Distributions of insertions identified in intergenic regions, introns, and exons. **(i)** GO-term enrichment analysis of closest genes to the insertions detected.

**Supplementary Fig. 9.**
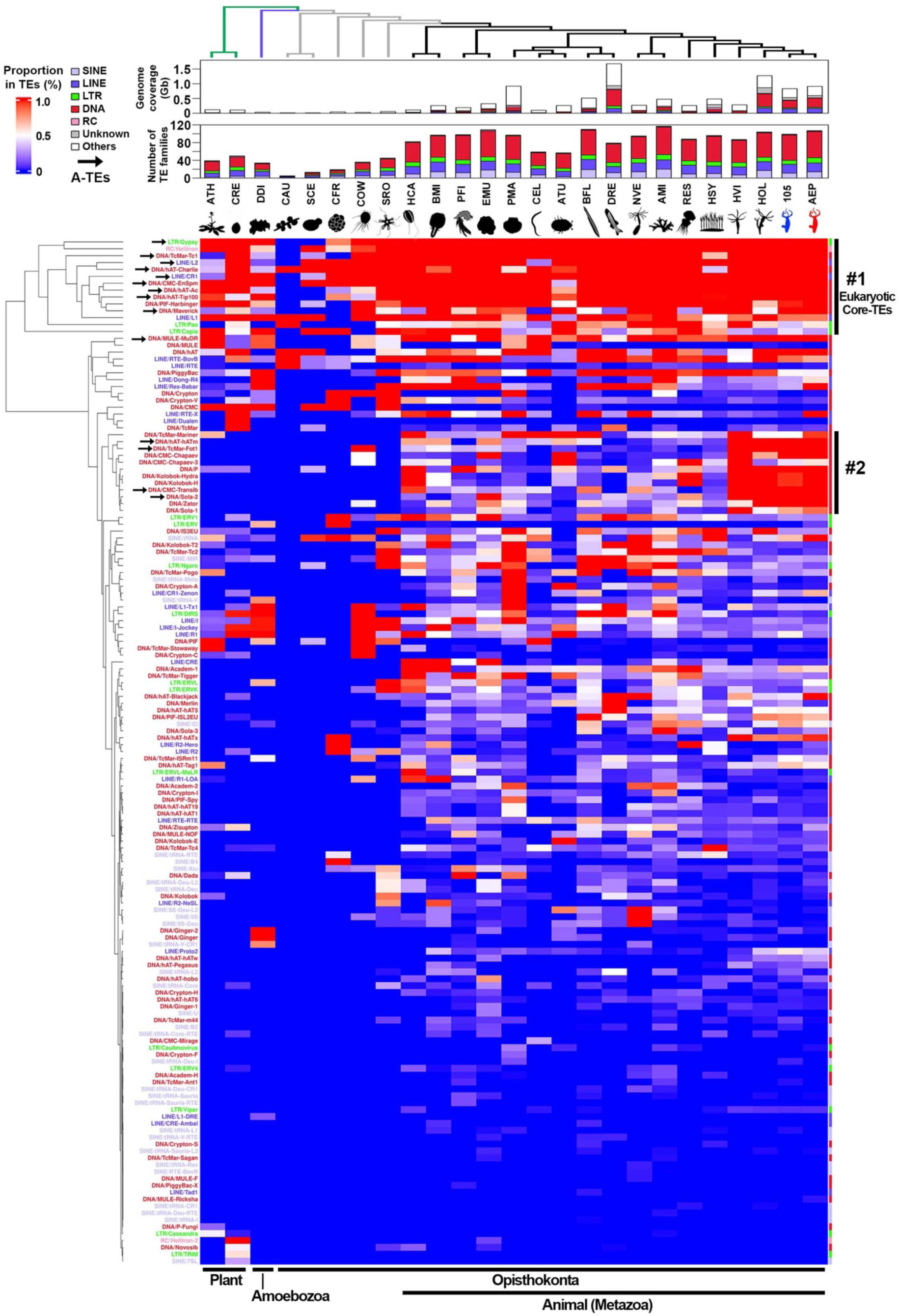
Deep TE homology across eukaryotes (related to. Fig. 5) Compositions of TEs in 17 metazoan genomes and those of eight outgroup species are shown. The heatmap represents the extent to which the genome coverage of each TE family constitutes the overall genome coverage of TEs for each species. The dendrogram at the left of the heatmap represents the result of hierarchical clustering performed using the Euclidean distance metric and the Ward’s method. Total 149 TE families detected in at least one of the 25 species analyzed are shown. Arrows indicate the 14 TEs belonging to the A-TEs. The cluster #1 and #2 are highlighted in Fig. 5. Abbreviations: CAU, *Candida auris*; SCE, *Saccharomyces cerevisiae*; CFR, *Creolimax fragrantissima*; COW, *Capsaspora owczarzaki*; SRO, *Salpingoeca rosetta*; HCA, *Hormiphora californensis*; BMI, *Bolinopsis microptera*; PFI, *Petrosia ficiformis*; EMU, *Ephydatia muelleri*; PMA, *Pecten maximus*; CEL, *Caenorhabditis elegans*; ATU, *Aethina tumida*; BFL, *Branchiostoma floridae*; DRE, *Danio rerio*; NVE, *Nematostella vectensis*; AMI, *Acropora millepora*; RES, *Rhopilema esculentum*; HSY, *Hydractinia symbiolongicarpus*; HVI, *Hydra viridissima*; HOL, *Hydra oligactis*.

**Supplementary Data 1.** Statistics of the genome assemblies

**Supplementary Data 2.** Gene annotation statistics

**Supplementary Data 3.** GeneBank accession numbers of ITS1-5.8S-ITS2 region of rDNA used for the phylogenetic analysis

**Supplementary Data 4.** Genome coverages of the TEs in the AEP genome and the 105 genome

**Supplementary Data 5.** TE families with cell type-specific expression

**Supplementary Data 6.** TE subfamilies with cell type-specific expression

**Supplementary Data 7.** TE loci with cell type-specific expression

**Supplementary Data 8.** Genome coverages of TEs in eukaryotes

## Notes

### Competing Interest Statement

The authors have declared no competing interest.

